# Identification of polymorphic and off-target probe binding sites on the Illumina Infinium MethylationEPIC BeadChip

**DOI:** 10.1101/056937

**Authors:** Daniel L. McCartney, Rosie M. Walker, Stewart W. Morris, Andrew M. McIntosh, David J. Porteous, Kathryn L. Evans

## Abstract

Genome-wide analysis of DNA methylation has now become a relatively inexpensive technique thanks to array-based methylation profiling technologies. The recently developed Illumina Infinium MethylationEPIC BeadChip interrogates methylation at over 850,000 sites across the human genome, covering 99% of RefSeq genes. This array supersedes the widely used Infinium HumanMethylation450 BeadChip, which has permitted insights into the relationship between DNA methylation and a wide range of conditions and traits. Previous research has identified issues with certain probes on both the HumanMethylation450 BeadChip and its predecessor, the Infinium HumanMethylation27 BeadChip, which were predicted to affect array performance. These issues concerned probe-binding specificity and the presence of polymorphisms at target sites. Using *in silico* methods, we have identified probes on the Infinium MethylationEPIC BeadChip that are predicted to (i) measure methylation at polymorphic sites and (ii) hybridise to multiple genomic regions. We intend these resources to be used for quality control procedures when analysing data derived from this platform.

## 1. Introduction

DNA methylation is an epigenetic mark typically occurring at cytosine-guanine dinucleotides (CpGs). Changes in DNA methylation are observed in normal development, in response to environmental stimuli, and in certain disease states [1]. DNA methylation is linked to transcriptional activity, rendering it a key regulatory motif [2]. Recent years have seen the development of high-throughput DNA methylation profiling techniques including whole-genome bisulphite sequencing (WGBS), methylated DNA immunoprecipitation (meDIP) and microarray-based technologies [3]. The Infinium HumanMethylation450 BeadChip, developed by Illumina, has offered an attractive array-based option to researchers, as it interrogates methylation at over 485,000 sites across the genome at single-base resolution at a relatively low cost (Bibikova et al., 2011 [4]). However, issues with probe-binding specificity and polymorphic targets have been identified which may compromise data integrity if not adequately addressed (Chen et al, 2013 [5]).

The Infinium HumanMethylation450 BeadChip has recently been superseded by the Infinium MethylationEPIC BeadChip. This array interrogates DNA methylation at over 850,000 sites, including > 90 % of the HumanMethylation450 array’s targets. This substantial increase in coverage, coupled with a continuing trend for interest in the role of DNA methylation, is likely to result in widespread use of this array. As such, it is essential that its potential shortcomings are thoroughly understood. In order to generate a resource that will be of use to researchers using the MethylationEPIC BeadChip we have identified probes that may perform sub-optimally. This work, therefore, represents an update of Chen et al.’s [5] previous characterisation of the Infinium HumanMethylation450 BeadChip.

Like its predecessor, the MethylationEPIC BeadChip uses two types of probe chemistry (Type I and Type II) to interrogate methylation. The differences between the two chemistries and the situations in which they are used have been described fully in previous publications [6]. Briefly, Type I assays use separate probes for unmethylated and methylated target sites while Type II assays use a single probe. Both assay types differentiate methylation state via single base extension of a fluorescent-labelled nucleotide.

Taking the differences between Type I and Type II assays into consideration, we have performed *in silico* analyses to identify probes on the Infinium MethylationEPIC BeadChip that are predicted to hybridise to multiple genomic regions, as well as probes where signal may be affected by polymorphisms at the target site, which could alter probe binding. Both of these factors should be taken into account when performing quality control of data produced using this technology.

## 2. Methods

### 2.1 Identification of probes with a polymorphic target

Probes potentially affected by polymorphisms at the target site were identified following methods described previously [5].

The signal-generating process of single-base extension requires end-nucleotide matching for both Type I and Type II probes. Therefore, we limited our query to target CpGs and sites of single-base extension, as polymorphisms at these sites are most likely to generate spurious signals.

Using information from the Infinium MethylationEPIC BeadChip manifest file (MethylationEPIC_v-1-0_B1.csv; date of download: 8 February 2016), we generated a list of genomic coordinates (hg19, GRCh37) of the target cytosine base (C) and guanine base (G) for all probes on the array. For Infinium Type I probes we also included the base before the target CpG, as this is the site of single base extension for these probes. We cross-referenced these coordinates to those of variants listed by the 1000 Genomes Project (phase 3) [7] to generate a list of probes affected by polymorphisms at the target CpG and/or site of single-base extension.

### 2.2 Identification of probes with non-specific hybridisation potential

Probes with the potential to cross-hybridise were identified following methods described previously [5].

#### 2.2.1 Generation of probe sequences for in silico analyses

Many Infinium Type II probe sequences contain an “R” nucleotide representing either an adenine (A) or guanine (G) base, depending on whether the underlying target cytosine is methylated or unmethylated. All possible combinations of Type II probe sequences were generated, and combined with a list of the Type I probe sequences.

#### 2.2.2 Generation of genomic comparison sequences for in silico analyses

The GRCh37 release of the human genome sequence was downloaded from the University of California, Santa Cruz (UCSC) Genome Browser website (https://genome.ucsc.edu/) as a reference, excluding alternative assemblies (e.g. chr17_ctg5_hap1) to avoid duplicated results (date of download: 11 January 2016). From this, we generated four modified reference genome sequences. A bisulphite-converted methylated forward genome sequence was generated *in silico* by converting all non-CpG cytosine bases to thymine (T) bases in the reference sequence. The same process was performed for the reverse complement of the reference sequence to generate a bisulphite-converted methylated reverse sequence of the human genome. Bisulphite-converted unmethylated forward and reverse sequences were generated by converting all C bases to T in the forward reference sequence and its reverse complement.

Using the BLAST-like alignment tool (BLAT) [8], we aligned the probe sequences described above to the four modified reference genome sequences, as well as their reverse complements. The BLAT parameters used were: *stepSize* = 5, *minScore* = 0, *minIdentity* = 0 and *repMatch* = 1,000,000,000. Probes were defined as being at high-risk of non-specific binding if there was a gap-free match of 47 or more nucleotides, which had to include the end base of the query sequence, at an off-target locus.

## 3. Results

### 3.1 Infinium MethylationEPIC BeadChip probes with polymorphic targets

Coordinates for 866,836 probes were obtained from the Infinium MethylationEPIC BeadChip manifest downloaded on 8^th^ February 2016. Excluding control probes, the manifest file contained 142,262 Type I probes (426,786 potential signal-affecting positions), and 724,574 Type II probes (1,449,148 potential signal-affecting positions), giving a total of 1,875,934 sites which were interrogated for genetic variation.

We identified 340,327 sites with either single nucleotide polymorphisms (SNPs), insertions or deletions (indels), or structural variation. These sites were targeted by 297,744 unique probes: 34% of the total probe content of the Infinium MethylationEPIC BeadChip. Of these, 23,399 probes (2.7%) targeted polymorphic sites with a minor allele frequency (MAF) of ≥ 5% in at least one population studied. A table of probes affected by polymorphisms, with minor allele frequencies corresponding to African, admixed American, European, South Asian, and East Asian populations (AFR, AMR, EUR, SAS, EAS; respectively) is available in the supplementary information of this paper (Supplementary Table 1).

### 3.2 Infinium MethylationEPIC BeadChip probes with cross-hybridisation potential

A total of 1,752,932 potential probe sequences, each 50 bases in length, were aligned to *in silico* bisulphite-converted forward and reverse methylated and unmethylated reference genomes, and their corresponding complementary strands in BLAT (i.e. eight single-stranded genomes in total). We identified 44,210 probes (11,772 Type I probes and 32,438 Type II probes) with ≥ 47 nucleotide off-target matches including the end base, which were defined as potentially cross-hybridising. A list of these probes is available in the supplementary information of this paper (Supplementary Tables 2-3).

Consistent with findings on the Infinium HumanMethylation450 BeadChip [5], a larger proportion of non-CpG-targeting probes (Probe ID prefix = “ch”) were identified as potentially cross-hybridising compared to CpG-targeting probes (Probe ID prefix = “cg”). Of 863,904 CpG-targeting probes present on the array, 42,558 (4.9% of total CpG-targeting probes) were identified as potentially cross-hybridising (Supplementary Table 2). In contrast, of 2,932 non-CpG targeting probes, we found only 1,280 to bind specifically to their targets while the remaining 1,652 were potentially cross-hybridising (56% of total non-CpG targeting probes; Supplementary Table 3), based on the information provided in the Illumina manifest.

## 4. Discussion

In order to identify probes that might compromise the performance of the Illumina Infinium MethylationEPIC BeadChip, we have generated lists of probes that may be affected by non-specific binding and/or polymorphisms at the target site.

Our *in silico* analyses identified 44,210 probes (5.1% total probe content) with potential off-target binding sites and 23,399 probes (2.7% total probe content) whose target site contains a polymorphism with a MAF ≥ 0.05 in at least one population studied, which may lead to artefactual signal due to impaired probe-binding. We recommend that users take these probes into consideration when analysing data on this platform, applying the appropriate filtering criteria in a population-specific manner, where possible. We recognise that there may be some situations where retaining probes mapping to polymorphic target sites will be desirable. For example, a difference in methylation due to a SNP that creates or destroys a CpG at a target site may be informative if it confers a change in disease risk.

Chen et al. (2013) [5] demonstrated that autosomal probes defined as potentially cross-hybridising according to their criterion of an off-target match of 47/50 bases, including the end nucleotide, showed an enrichment for off-target binding sites on the sex chromosomes. Failure to exclude these probes could, therefore, result in the spurious conclusion that these loci are differentially methylated between males and females. Following their methods, we have identified probes on the Infinium MethylationEPIC BeadChip with the potential to hybridise to multiple genomic regions, thus generating off-target signal. We suggest the exclusion of these probes prior to data analysis. Although the exclusion of potentially cross-hybridising probes defined using this method is likely to result in an improvement in the validity of the results obtained from the array, it is likely that the actual extent of off-target binding will vary by locus. Factors such as local sequence composition, including the presence of polymorphisms underlying the probe sequence, are likely to play a role in determining the likelihood of cross-hybridisation. It is, therefore, recommended that any results of interest that may have been generated due to cross-hybridisation are checked using an alternative technique, such as pyrosequencing of bisulphite-converted DNA.

In summary, we have produced lists of probes on the new Illumina Infinium MethylationEPIC BeadChip that measure methylation at sites affected by polymorphisms and/or have the potential to cross-hybridise. Based on the wide-spread use of the HumanMethylation450 BeadChip, we predict that the Illumina Infinium MethylationEPIC BeadChip will play a central role in epigenome-wide association studies (EWAS) over the next few years. As such, it is essential that factors affecting the performance of the array, such as probe specificity and sequence polymorphisms, which we have demonstrated to potentially affect a substantial proportion of probes, are taken into consideration. We recommend that the resources supplied with this paper be used in conjunction with additional standard quality control measures, such as excluding probes with low signal-to-background ratios, omission of samples with a high proportion of such probes, and appropriate data normalisation strategies (for review see Wilhelm-Benartzi et al., 2013 [9]), in order to maximise the likelihood of producing meaningful results.

## Acknowledgements

DLMcC is funded by a Ph.D. studentship awarded by Mental Health Research UK (http://www.mhruk.org). KLE would like to acknowledge the support of the Brain & Behavior Research Foundation through a NARSAD Independent Investigator Award. AMM would like to acknowledge the support of The Health Foundation through a Clinician Scientist Fellowship and a NARSAD Independent Investigator Award from the Brain & Behavior Research Foundation and Wellcome Trust Strategic Support (104036/Z/14/Z). KLE, AMM and DJP are members of The University of Edinburgh Centre for Cognitive Ageing and Cognitive Epidemiology (CCACE), part of the cross council Lifelong Health and Wellbeing Initiative (G0700704/84698). Funding of CCACE from the BBSRC, EPSRC, ESRC and MRC is gratefully acknowledged.

